# Deamidated TPI is an efficacious target for cell-selective therapy in triple-negative breast cancer

**DOI:** 10.1101/2021.11.09.467888

**Authors:** Sergio Enríquez-Flores, Luis A. Flores- López, Ignacio De la Mora-De la Mora, Itzhel García-Torres, Isabel Gracia-Mora, Pedro Gutiérrez-Castrellón, Cynthia Fernández-Lainez, Yoalli Martínez-Pérez, Alberto Olaya-Vargas, Paul de Vos, Gabriel López-Velázquez

**Affiliations:** Laboratorio de Biomoléculas y Salud Infantil, Instituto Nacional de Pediatría, CDMX, México; CONACYT Instituto Nacional de Pediatría, CDMX, México; Directora de la Unidad de Investigación Preclínica, Facultad de Química, Universidad Nacional Autónoma de México, CDMX, México; Hospital General Dr. Manuel Gea González, México; Laboratorio de Errores Innatos del Metabolismo y Tamiz, Instituto Nacional de Pediatría, CDMX, México; Stem Cell Transplantation and Cellular Therapy, Instituto Nacional de Pediatría, CDMX, México; Department of Pathology and Medical Biology, University of Groningen, University Medical Center Groningen, Groningen, 9713 GZ, the Netherlands; Posgrado en Ciencias Biológicas, Universidad Nacional Autónoma de México, CDMX, México

**Keywords:** cancer, drug repurposing, drug target, glycolysis, methyl glyoxal, triosephosphate isomerase

## Abstract

Human TPI (HsTPI) is a central and essential glycolytic enzyme for energy supply and is overexpressed in cancer cells. Here, we investigated HsTPI as a potential target for inducing cell death in triple-hormone receptor-negative breast cancer, which is highly dependent on glycolysis, and therapies for its treatment are limited. We found endogenous accumulation of deamidated HsTPI in human breast cancer cells, which might be caused by the lower activity of the HsTPI-degrading caspase-1 in breast cancer cells. *In silico* and *in vitro* analyses of deamidated HsTPI demonstrated the efficacy of thiol-reactive drugs in blocking enzyme activity. The cancer cells were selectively programmed to undergo apoptosis with thiol-reactive drugs by inducing the production of methylglyoxal (MGO) and advanced glycation-end products (AGEs). *In vivo* in mice, the thiol-reactive drug effectively inhibited the growth of human tumors by targeting HsTPI as underlying mechanism. Our findings demonstrate deamidated HsTPI as a novel target to develop therapeutic strategies for treating cancers and other pathologies in which this post-translationally modified protein accumulates.

---

Since cancer cells show a high capacity for proliferation, they are energy demanding which is driven by glycolysis[1-3]. Due to this high glycolysis and upregulation of glycolysis-associated enzymes in cancer cells[4-6], substrate analogs for glycolytic enzymes have been proposed as potential anticarcinogenic therapeutics[7-9] Triosephosphate isomerase (TPI or TIM) is such a potential target for these drugs as it plays a central role in the energy generating phase of glycolysis by isomerizing dihydroxyacetone phosphate (DHAP) to D-glyceraldehyde-3-phosphate (GAP). Regions distinct from catalytic sites are promising for efficacious anti-tumor strategies. These arguments were the rational to investigate the use of deamidated human TPI (HsTPI) as a druggable target[10].

In eukaryotes, deamidation is a spontaneous modification of proteins that has repercussions for the proteins’ activity and stability, both *in vivo* and *in vitro*[11-13]. Especially, asparagine (Asn) and glutamine (Gln) are targets for deamidation in tumor cells. Asparagine deamidation into aspartic acid (Asp) and isoAsp results in alteration of the primary structure of the protein, introducing a negative charge, and often modifying their tertiary structure. As a consequence, functional aspects of the protein are lost interfering with stability of the cells.

Two deamidation sites in HsTPI at positions 16 and 72 have been found. Deamidation of Asn 16 is the most common and induces major functional and structural alterations of HsTPI [12]. Continuous catalytic cycles promote the deamidation of HsTPI [14], which is boosted in aged cells as well as in cancer cells [15]. Studies on the structural characteristics of deamidated HsTPI have demonstrated that it is prone to selective inhibition with thiol-reactive compounds [10] and therefore be an effective approach to target HsTPI and there with interfering with glycolysis in cancer cells.

Since glycolysis is not adequately regulated in cancer cells, we hypothesized that deamidated HsTPI commonly accumulates in these cells, enabling its use as an efficient druggable target to selectively exert cell death. Here, the presence and accumulation of deamidated HsTPI were demonstrated in triple-hormone receptor negative breast cancer cells, which showed high sensitivity to the thiol-reactive drugs rabeprazole and auranofin. Both drugs inhibited cellular HsTPI enzyme activity and induced selective cell death. Nude immunocompromised mice with implanted human breast cancer cells treated with rabeprazole exhibited significantly reduced tumor size compared with the tumor size in untreated mice. Additionally, pretreatment of cancer cells with rabeprazole did not lead to tumor formation after implantation in mice. The central finding of our study is that deamidated HsTPI is naturally present and accumulates in cancer cells but not in noncancer cells. Overall, the results of *in vitro* (recombinant protein), *in situ* (cell culture) and *in vivo* (xenograft murine model) experiments demonstrate that deamidated HsTPI is an efficient druggable target.

## RESULTS

### Deamidated HsTPI is highly inactivated and structurally blocked by the drugs rabeprazole and auranofin through Cys modification

HsTPI is a homodimeric enzyme for which deamidation occurs at amino acid residues located in the contact site between its two subunits (interface), increasing its accessibility for small ligands[10, 12]. To gain more insight in its active ligand binding sites, we performed *in silico* and *in vitro* analyses of crystallographic structures of nondeamidated and deamidated HsTPI and the corresponding recombinant proteins. The largest and deepest cavity at the interface was found in deamidated HsTPI. Docking of rabeprazole and auranofin identified more binding sites in deamidated HsTPI than in nondeamidated HsTPI (Fig. 1a).

**Fig. 1.**
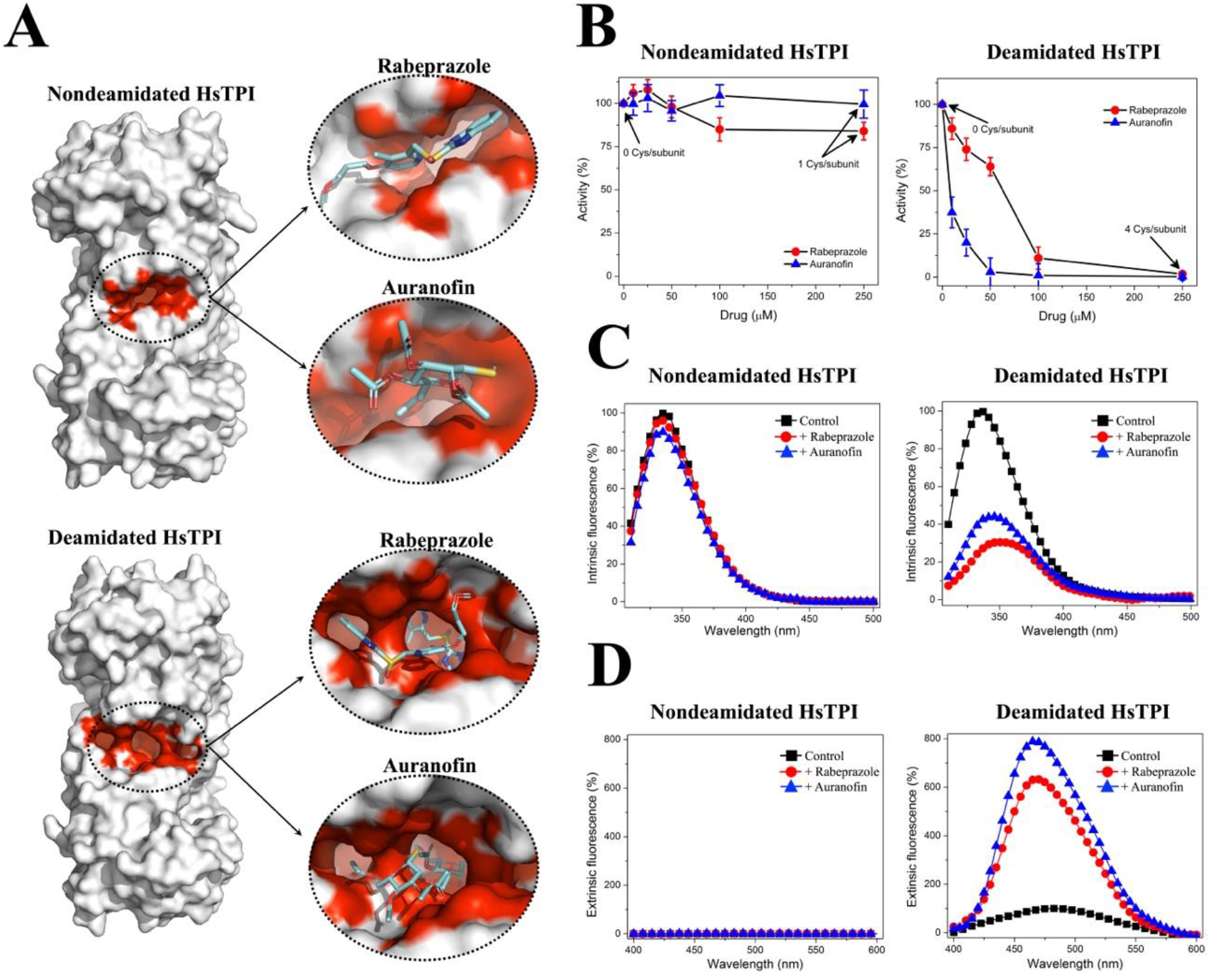
Deamidated HsTPI was more likely to be functionally and structurally disturbed by the thiol-reactive drugs rabeprazole and auranofin than its nondeamidated counterpart. **a**, Ligand docking in nondeamidated and deamidated enzymes. Rabeprazole and auranofin were docked at the interface of the crystallographic structure. As shown, the deamidated protein incorporates more ligands in the innermost part of the interface than its nondeamidated counterpart. **b**, Inactivation assays with recombinant nondeamidated and deamidated HsTPI proteins, with rabeprazole 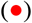 or auranofin 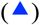. After incubation, the residual enzymatic activity was determined. The results are expressed as the percent activity, with the enzyme activity without the drug set at 100%. **c**, Intrinsic fluorescence of nondeamidated and deamidated HsTPI with or without drug treatment, in the absence 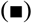 (control) or presence of 250 µM rabeprazole (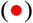 or 250 µM auranofin 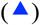. The results are expressed as the percent fluorescence, with the maximum fluorescence emission of the control enzyme set at 100%. **d**, Extrinsic fluorescence of ANS corresponding to the enzymes with or without drug treatment in the absence 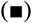 (control) or presence of 250 µM rabeprazole (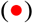 or 250 µM auranofin 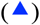. After incubation, the extrinsic fluorescence was determined by adding 100 µM ANS. The background was always subtracted. The results are expressed as percent of fluorescence, with the maximum fluorescence emission of the control in deamidated HsTPI set at 100%. The results represent the mean of four independent experiments.

Consistent with the docking data, our functional *in vitro* studies with human breast cancer cells showed that in contrast to nondeamidated HsTPI, its deamidated counterpart was selectively bound by both drugs, which completely inhibited its enzyme activity (Fig. 1b, right). Additionally, inhibition of deamidated HsTPI was dose dependent, and deamidated HsTPI was 4-fold more sensitive to rabeprazole and 30-fold more sensitive to auranofin than nondeamidated HsTPI.

The main mechanism of action of both drugs (rabeprazole and auranofin) is covalent binding to the thiolate moieties of cysteine residues (Cys) (named Cys derivatization) [16, 17]. The primary structure of HsTPI contains 5 Cys residues per subunit, which were visualized and quantified in the recombinant enzyme in absence of the drugs. Treatment with either rabeprazole or auranofin derivatized 4 Cys residues per subunit in deamidated HsTPI, whereas nondeamidated HsTPI contained only one derivatized Cys residue per subunit after drug treatment (Fig. 1b, and Suppl. Table 1).

In addition to these functional studies, we carried out spectroscopic studies to identify structural changes due to drug treatment. To that end, we analyzed the intrinsic fluorescence of HsTPI to follow changes in the exposure of its aromatic residues to solvent in presence of the drugs. The intrinsic fluorescence spectra of nondeamidated HsTPI showed marginal changes in the tertiary structure of the enzyme upon treatment with either rabeprazole or auranofin (Fig. 1c, left). In contrast, the intrinsic fluorescence spectra of deamidated HsTPI upon treatment with both drugs decreased by more than 50% (Fig. 1c, right). Under these conditions, rabeprazole and auranofin generated a 70% and 56% decrease in fluorescence intensity and shifts in the maximum fluorescence emission (λ_max_) of 12 and 9.5 nm, respectively (Fig. 1c, right). A shift of λ_max_ toward a longer wavelength (redshift) indicates exposure to tryptophan residues previously buried into a folded structure.

To substantiate proof for the structural changes of deamidated HsTPI and the drugs, extrinsic fluorescence studies with 8-anilinonaphthalene-1-sulfonic acid (ANS) were performed. ANS is a fluorescent molecular probe that has a high affinity for hydrophobic regions; thus, proteins containing more hydrophobic cavities will exhibit higher ANS fluorescence signals. The ANS fluorescence in the untreated and treated nondeamidated HsTPI showed a marginal signal (Fig. 1d, left). In contrast, the fluorescence signal from ANS in deamidated HsTPI was strongly increased under all conditions. This signal was augmented with respect to that of the untreated enzyme by 6.5- and 8-fold upon treatment with rabeprazole and auranofin, respectively (Fig. 1d, right). Additionally, the λ_max_ shifted by 14 nm to a lower wavelength (shift to blue) with both drugs. These results show that as a consequence of the drug treatments, deamidated HsTPI became highly accessible to small molecules such as ANS.

### Deamidated HsTPI is present and accumulates in breast cancer cells

To study possible endogenous accumulation of deamidated HsTPI in breast cancer cells, a previously validated method of selective cleavage by hydroxylamine was used[18, 19]. Hydroxylamine can selectively cleave the intermediary succinimide that is formed by the side chains of Asn and glycine (Gly) in nondeamidated proteins under alkaline conditions; furthermore, in deamidated proteins, a chain composed of Asp-Gly (or isoAsp-Gly) does not form succinimide under the same conditions. Therefore, the deamidated protein is not cleaved[20]. Then, as HsTPI contains two Asn-Gly pairs at positions 16-17 and 72-73, cleavage might generate three peptides that are 1.87, 5.96 and 18.87 kDa in size if the enzyme is not deamidated. When the cleavage of nondeamidated HsTPI is incomplete, a peptide with a size of 7.83 kDa could also be produced. On the other hand, the cleavage of deamidated HsTPI at only position 16 would produce peptides with sizes of 7.83 and 18.87 kDa, whereas double deamidated HsTPI would not be cleaved at position 16 or 72. This difference in cleavage patterns of the recombinant enzymes *versus* cellular HsTPI from human primary mammary epithelial cells (HMECs) (normal breast cells) and MDA-MB-231 cells (triple-negative breast cancer cells) was used to study differences in accumulation of deamidated HsTPI.

HsTPI from HMECs showed the same cleavage pattern found for recombinant nondeamidated HsTPI (Fig. 2a, lane 6 *vs* lane 3), whereas HsTPI from MDA-MB-231 cells showed a cleavage pattern similar to that of recombinant HsTPI deamidated at position 16 (Fig. 2a, lane 7 *vs* lane 4). The cleavage selectivity of the hydroxylamine method was reinforced by the absence of a cleavage pattern for the recombinant double deamidated HsTPI at positions 16 and 72 (Fig. 2a, lane 5). Herewith we demonstrated that deamidated HsTPI was present in these cancer cells and not in their noncancerous counterparts.

**Fig. 2.**
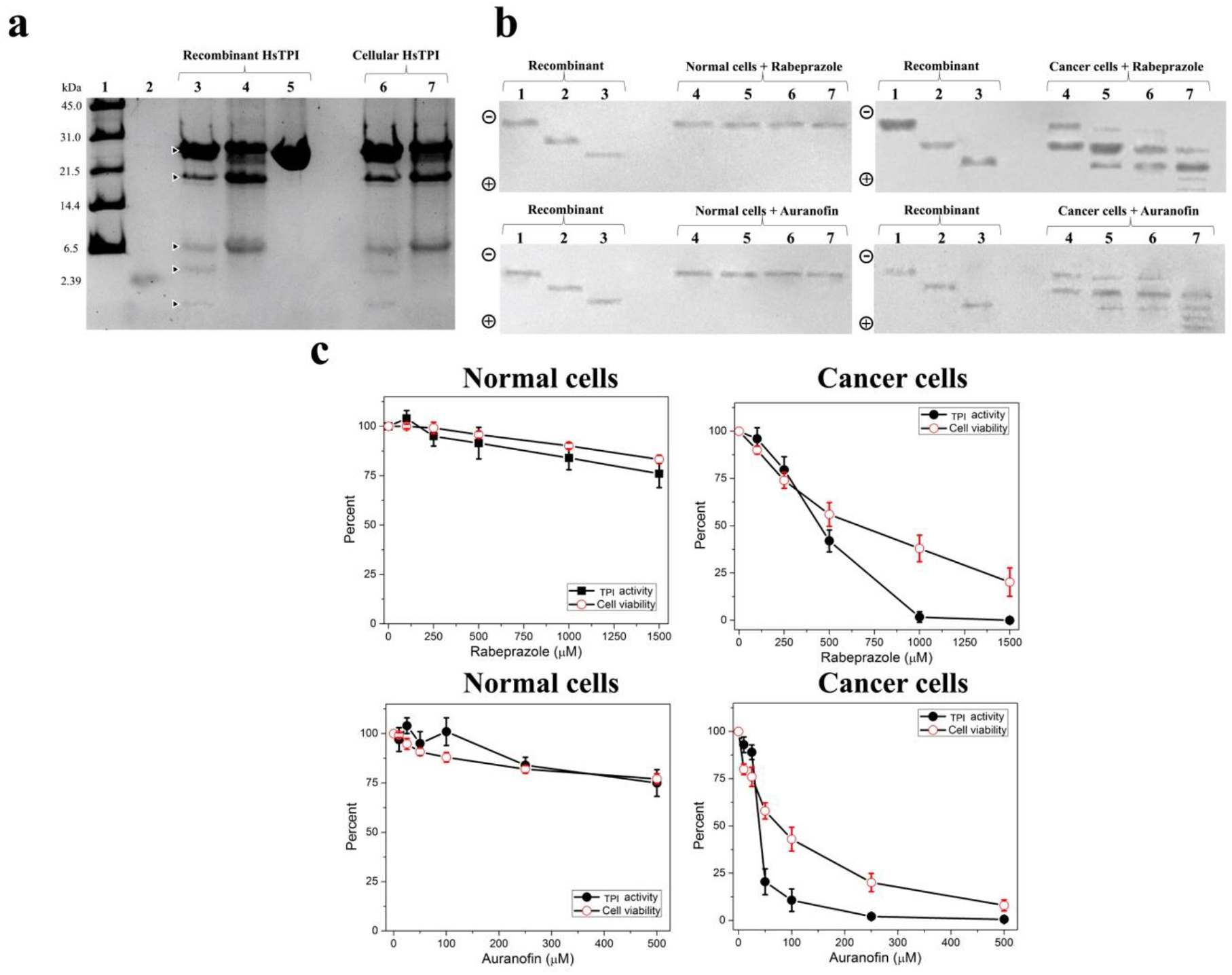
Deamidated HsTPI accumulated in breast cancer cells was affected by treatment with rabeprazole and auranofin. **a**, Hydroxylamine cleavage of the recombinant and cellular HsTPI proteins. Lane 1: molecular weight standard (broad range, Bio-Rad), lane 2: peptide with a size of 2.39 kDa, lanes 3 to 5: (10 µg/lane) nondeamidated (WT), once deamidated (N16D) and double deamidated (N16D/N72D) HsTPI proteins. Lanes 6 and 7: (20 µg/lane) immunoprecipitated HsTPI from HMECs and MDA-MB-231 cells, respectively. The cleavage pattern of the nondeamidated recombinant enzyme (lane 3) is similar to that of cellular HsTPI from normal cells (HMECs) (lane 6). In the same manner, the cleavage pattern of the deamidated recombinant enzyme (lane 4) is similar to that of cellular HsTPI from cancer cells (MDA-MB-231) (lane 7). **b**, Western blot analysis of recombinant and cellular HsTPI in nPAGE. In the four panels, lanes 1 to 3 (10 µg of protein/lane) correspond to the recombinant nondeamidated, once deamidated and double deamidated HsTPI, respectively. Lanes 4 to 7 (100 µg protein/lane) correspond to proteins from the protein extract from normal cells (HMECs) (left panel, upper and bottom) and cancer cells (MDA-MB-231) (right panel, upper and bottom) treated with 0, 500, 1000 and 1500 µM rabeprazole (upper) or 0, 25, 50 and 100 µM auranofin (bottom). **c**, HsTPI enzyme activity and viability of HMECs and MDA-MB-231 cells treated with rabeprazole and auranofin. The results are expressed as percentages of residual enzyme activity, with the values obtained in the absence of the drugs set at 100%. The results are the mean of four independent experiments.

Since every deamidation reaction adds one negative charge to HsTPI, native polyacrylamide gel electrophoresis (nPAGE) can be applied for analyzing the acidic *status* of both recombinant and cellular HsTPI proteins. The electrophoretic mobility patterns resulting from this immunoblotting of the nPAGE gel clearly showed and confirmed that in cancer cells deamidated HsTPI accumulated (Fig. 2b, cancer cells, lane 4), which was not observed in normal cells (Fig. 2b, normal cells lanes 4). These observations were the main rational to propose and test drugs that target and inhibit deamidated HsTPI to selectively delete cancer cells and keep non-cancerous cells virtually untouched.

### Rabeprazole and auranofin induce the formation of inactive acidic isoforms of HsTPI in cancer cells and promoted selective cell death

Negatively charged HsTPI isoforms (acidic isoforms) appeared after the treatment of cancer cells (Fig. 2b, cancer cells, lanes 5-7) but not in normal cells (*i*.*e*. noncancer cells) (Fig. 2b, lanes 5-7) with rabeprazole or auranofin. Furthermore, the naturally accumulated deamidated HsTPI already present in the cancer cells tended to disappear, and other more negatively charged isoforms appeared as a result of treatment with increasing doses of both drugs. Importantly, both drug treatments induced the disappearance of nondeamidated HsTPI, which potentiates the effect of the drugs on cancer cells.

Cellular HsTPI in cancer cells completely lost its enzyme activity upon treatment with 1000 µM rabeprazole, and the viability of these cells decreased to approximately 80% at 1500 µM rabeprazole treatment (Fig. 2c, cancer cells, rabeprazole). Auranofin strongly reduced cellular HsTPI levels in cancer cells at lower concentration. With 250 µM auranofin enzyme activity was totally inhibited and cell viability was reduced to 80%. At a concentration of 500 µM auranofin totally eliminated cell viability (Fig. 2c, cancer cells, auranofin). These drugs exerted effects on the enzyme activity of cellular HsTPI similar to those obtained for the recombinant enzymes.

In contrast, in normal cells, the two drugs induced no more than a 25% loss in both enzyme activity and cell viability (Fig. 2c, normal cells, rabeprazole and auranofin) again illustrating its selectivity for cancerous cells.

Overall, these results show that HsTPI in cancer can be selectively targeted, supporting the crucial involvement of this protein in maintaining viability in breast cancer cells and the selective cell death that can be induced by the tested drugs.

To support the notion that the inhibition of cellular HsTPI was attributable to the drug treatments instead of the consequence of cell damage, assays with high drug concentrations and short incubation times were performed. HsTPI activity in breast cancer cells treated with rabeprazole was almost totally abolished at 2 h of treatment, and the viability of the cells was close to 100% for 4 h until it dropped by approximately 20% in the last hour (Suppl. Fig. 1, cancer cells, rabeprazole). With auranofin, HsTPI activity was completely abolished at 4 h, whereas cell viability remained close to 100% and tended to decrease after 5 h of treatment (Suppl. Fig. 1, cancer cells, auranofin). This effect was selective for cancerous cells as HsTPI activity in normal cells dropped by not more than 30% with rabeprazole treatment, while auranofin treatment exerted a maximum inhibitory effect of 10%. The viability of these cells was maintained close to 100% during treatment (Suppl. Fig. 1, normal cells).

These results again support the observation that TIM in cancer cells is a cell-specific target of both drugs that is inactivated prior to cell death, whereas TIM in normal cells is practically unaffected. To assess the hypothesis that the drugs direct their effect to regions other than the catalytic site, the kinetics of cellular HsTPI were determined for the latter assayed conditions in untreated and treated cells. The *K*_m_ values for TIM in untreated and treated cancer cells were similar; however, the V_max_ values were 3.3 (with rabeprazole) and 2.6 (with auranofin) times lower than those in untreated cells (Suppl. Fig. 2 and Suppl. Table 2). Consistent with previous reports[12, 21], the *K*_m_ values of TIM in normal cells, either untreated or treated with rabeprazole and auranofin, were similar (Suppl. Fig. 2, and Suppl. Table 2). These results support the idea that these drugs do not compete with the active site of HsTPI and that their mechanism of inactivation is noncompetitive.

### Targeting deamidated HsTPI decreased lactate production and induced excessive production of methylglyoxal and AGEs in breast cancer cells

Targeting HsTPI disturbs the glycolytic pathway, which can be demonstrated and quantifying by measuring lactate production. Breast cancer cells, which normally produce high quantities of lactate, significantly decreased their production of lactate when treated with either rabeprazole or auranofin, whereas normal cells were not affected (Fig. 3a). This supports the idea that, when HsTPI is affected by these drugs, glycolytic flux is depleted in cancer cells (Fig. 3a, cancer cells). Additionally, the malfunctioning of HsTPI causes the accumulation of DHAP, the spontaneous degradation of which leads to methylglyoxal (MGO) formation, which in turn inhibits glycolytic flux [22]. An increase in MGO is related to HsTPI failure, documented in several human disorders [23]. Concordantly, our results showed that the tested treatments increased the production of MGO, but such production was significantly higher in cancer cells than in normal somatic cells (Fig. 3b, cancer cells).

**Fig. 3.**
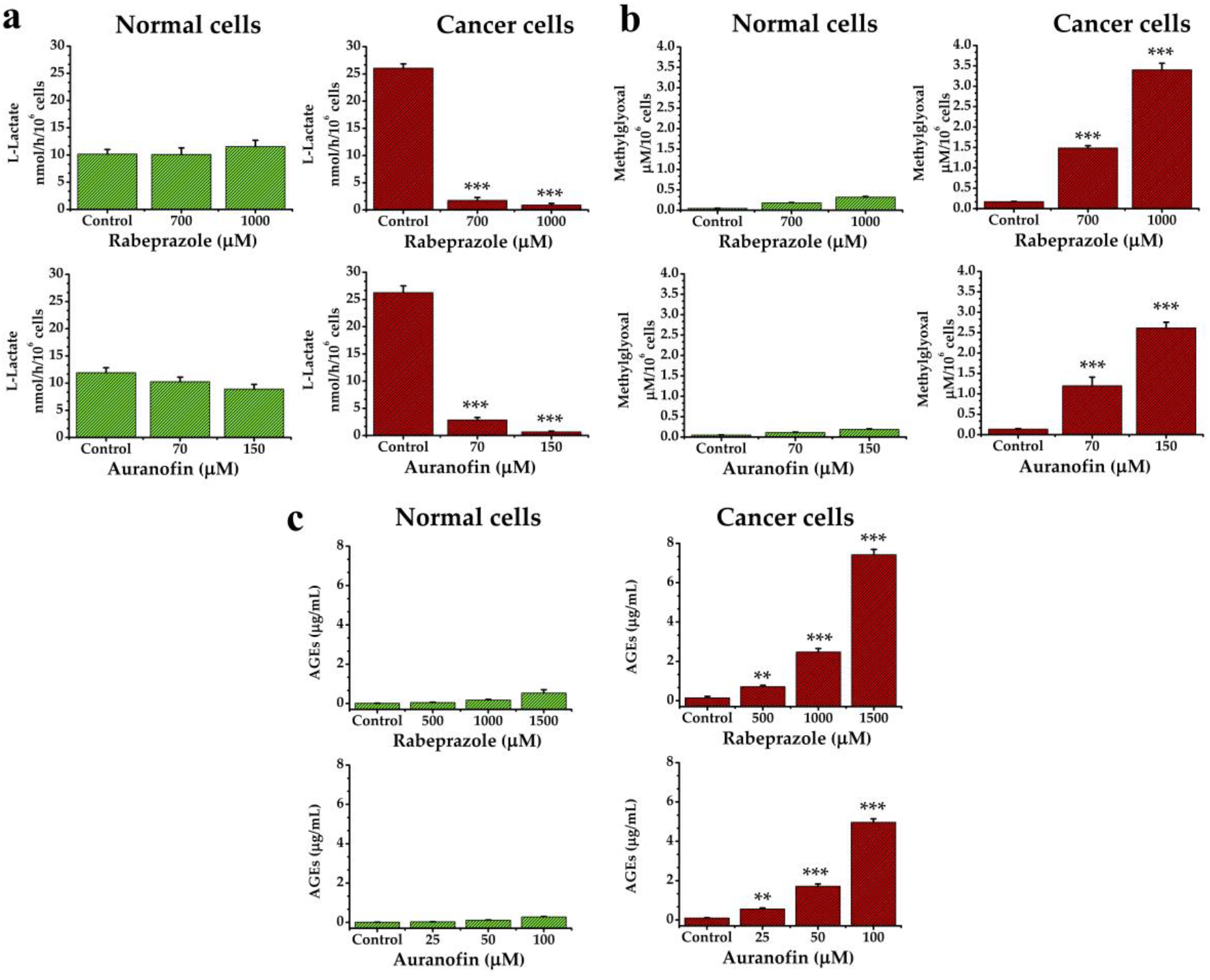
Quantification of metabolites in normal and cancer cells under drug treatments. **a**, Determination of extracellular lactate levels in normal and cancer cells. Cells were incubated with 0, 700, and 100 µM of rabeprazole or 0, 75, and 150 µM of auranofin. The extracellular lactate level is shown. **b**, Determination of MGO levels in normal and cancer cells with rabeprazole (0, 700, and 1000 µM) or auranofin (0, 75 and 150 µM). Intracellular MGO is shown. **c**, Determination of AGE levels in normal and cancer cells. with 0, 500, 1000 and 1500 µM rabeprazole or 0, 25, 50 and 100 µM auranofin. All results represent the mean of at least four independent experiments. Differences among groups were assessed with one-way ANOVA and Tukey’s test with *p* value = 0.001 *** or with *p* value = 0.01**

MGO is harmful to several cell types, mainly as it is an essential intermediate in the formation of AGEs. In this regard, both drug treatments significantly increased the levels of AGEs in a dose-dependent manner in cancer cells (Fig. 3c, cancer cells) but not in normal cells, in which these products were present at minimal concentrations (Fig. 3c, normal cells). These results are consistent with those observed for MGO formation upon the treatments. Hence, the behavior of deamidated HsTPI leads to the production of toxic metabolites (*i*.*e*., MGO and AGEs), which in turn should be involved in the selective cell death process that we observed.

### Breast cancer cells treated with rabeprazole and auranofin underwent apoptosis

Enhancing apoptosis is important to prevent tumor initiation, progression and metastasis formation [24]. As shown in our previous experiments, rabeprazole and auranofin induce overproduction of MGO and AGEs. Both metabolites are known to induce apoptosis [25, 26]. To test the influence of rabeprazole and auranofin on cell apoptosis, we analyzed the expression of proteins related to pro- and antiapoptotic events [27] induced through intrinsic pathways (Bax and Bcl-2) [28, 29].

The results showed that both drug treatments induced a decrease in the expression of total ERK 1/2 and Bcl-2. Additionally, phosphorylated ERK 1/2 levels were decreased (Fig. 4a). These elements are related to lower cellular proliferation. On the other hand, the proapoptotic elements Bax and cleaved procaspase-7 were increased after both treatments in a dose-dependent manner (Fig. 4a). Additionally, transferase dUTP nick end labeling (TUNEL) assays with breast cancer cells treated with rabeprazole or auranofin showed an increase in apoptotic cells after 24 h of treatment (Fig. 4b). These results show that our strategy overcomes the antiapoptotic tendencies of cancer cells.

**Fig. 4.**
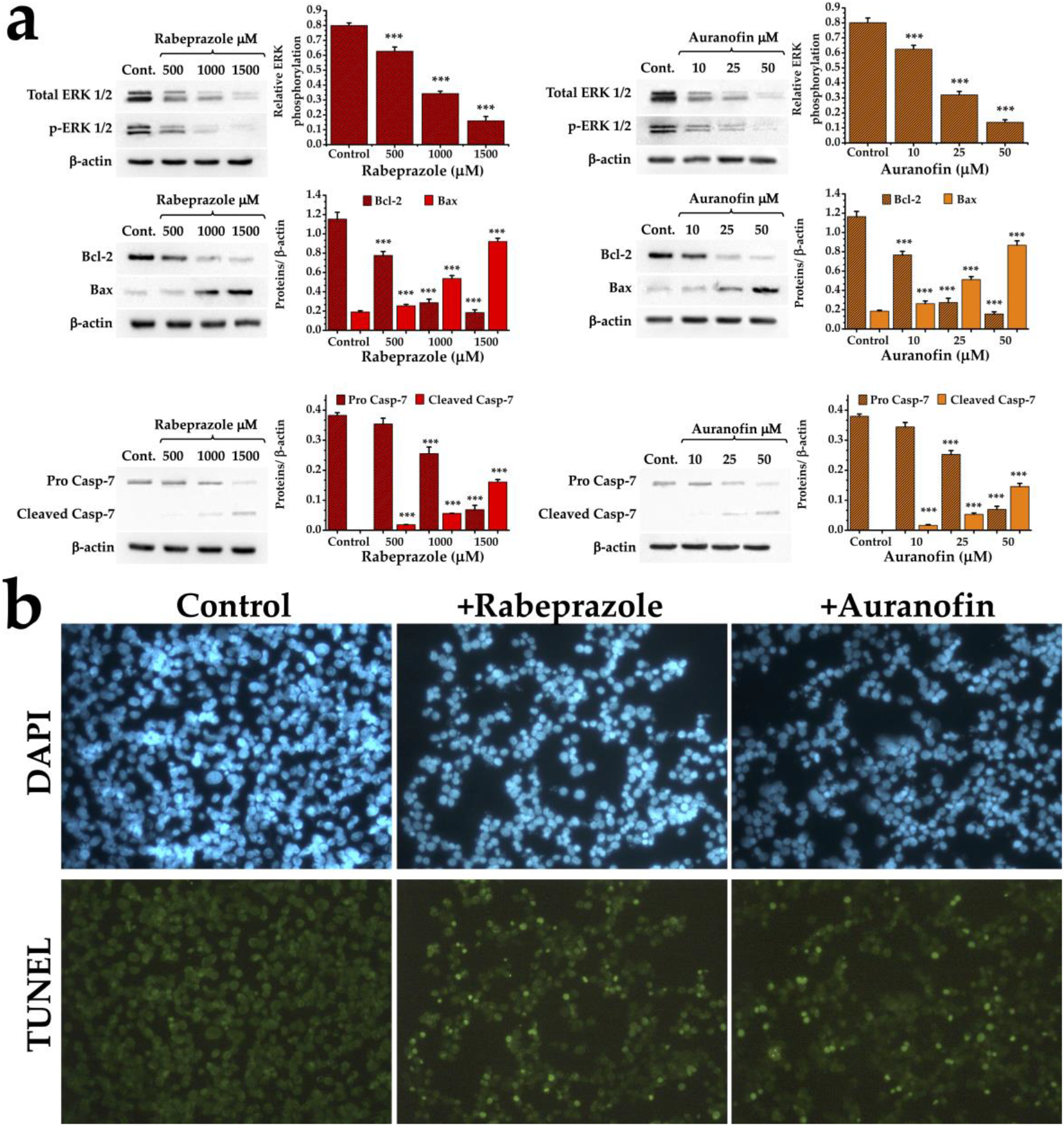
The selective cell death of cancer cells upon drug treatment occurs by means of apoptosis. **a**, Cancer cells were exposed to 0, 500, 1000 and 1500 µM rabeprazole and 0, 10, 25 and 50 µM auranofin. Expression levels of the ERK 1/2 pERK 1/2, Bcl2, Bax, procaspase-7 and caspase-7 proteins are shown. β-Actin was used as control. Graphs on the right side of the panels correspond to the results of densitometric analysis of each condition. **b**, TUNEL assays with cancer cells that were or were not treated with the drugs. Upper panels correspond to DAPI nuclei staining, and the bottom panels correspond to TUNEL assays. Photographs are at 40×.

### Deamidated HsTPI is naturally accumulated in tumoral breast cancer tissues and is efficient as a druggable target

To determine whether the properties of deamidated HsTPI observed *in vitro* would translate into an efficient anticarcinogenic strategy, human breast cancer cells were implanted in nude immunocompromised mice and treated with rabeprazole. Tumorigenesis was significantly inhibited by three times weekly treatment with rabeprazole (Fig. 5a) and totally inhibited when pretreated cancer cells were implanted (Fig. 5a, pretreatment). The most notable results from these series of experiments were a) the finding that deamidated HsTPI naturally accumulated in the tumorigenic cells and b) the identical electrophoretic mobility behaviors of tumorigenic HsTPI obtained *in vitro* and treated with rabeprazole combined with the high production of AGEs in tumors from treated mice (Fig. 5b).

**Fig. 5.**
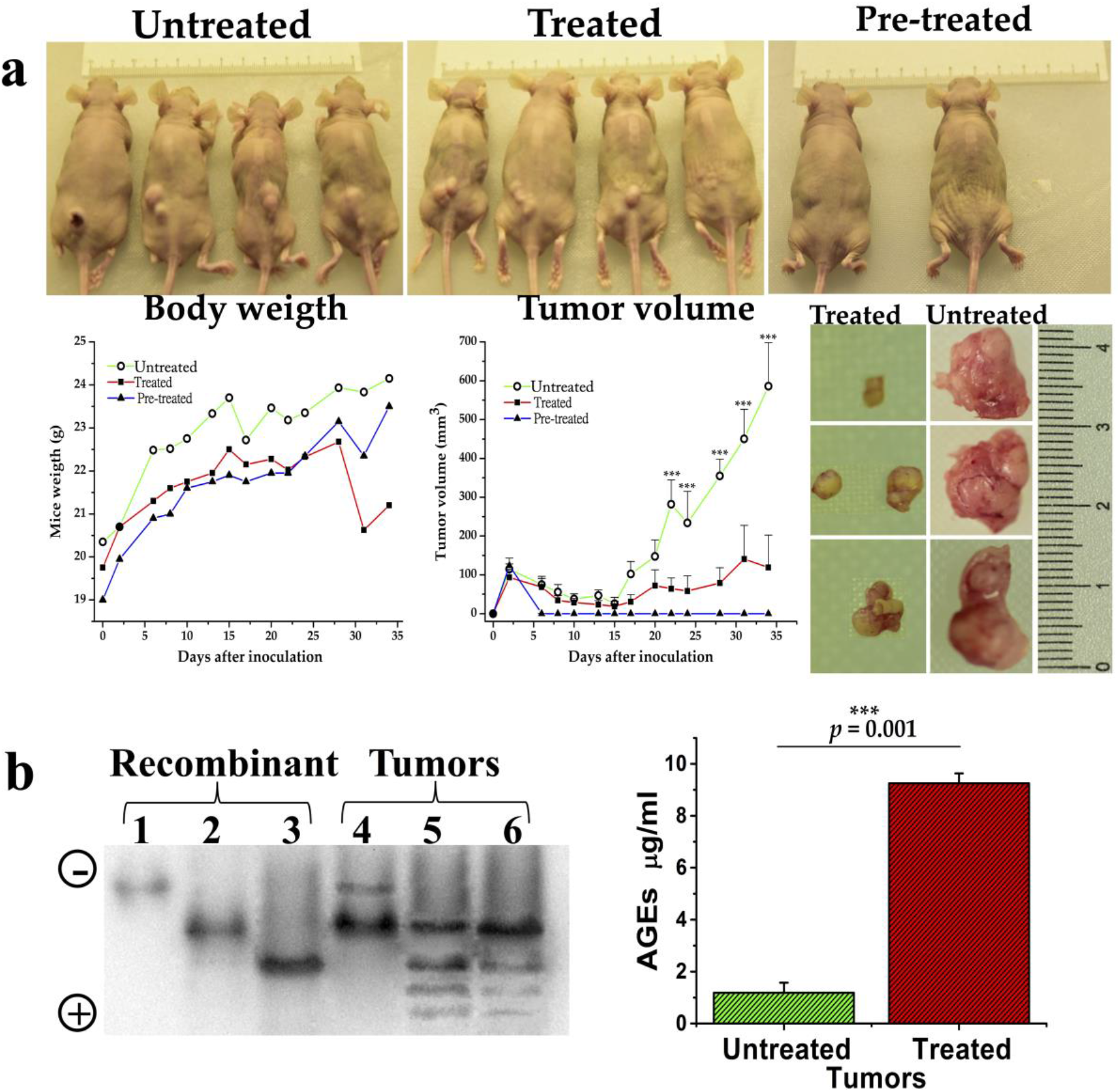
Deamidated HsTPI accumulated in xenograft tumors, and the antitumorigenic activity of rabeprazole was accompanied by change in HsTPI behavior and the overproduction of AGEs. **a**, Representative images of mice from the different experimental groups (upper panel). Changes in body weight and tumor volume were noted throughout the experiment (graphs, left bottom). Representative images of tumors (photographs, right bottom) from treated (left) and untreated (right) mice at the end of the experiment clearly depict the antitumorigenic effect of rabeprazole. Mice implanted with breast cancer cells previously incubated for 24 h with 1 mM rabeprazole (pretreated group) did not develop tumors (upper panel, right). Differences among groups were assessed with one-way ANOVA and Tukey’s test with *p* value = 0.001 ***. **b**, Acidic isoforms of HsTPI obtained from tumors and compared with the recombinant HsTPIs (left panel). Lanes 1-3 show the recombinant proteins and were used as patterns to identify the acidic isoforms corresponding to the nondeamidated (lane 1), once deamidated (lane 2), and twice deamidated HsTPI. Lanes 4-6 show the acidic isoforms of HsTPI from the xenograft tumors of untreated mice (lane 4) and the acidic isoforms of HsTPI from the xenograft tumors of treated mice (lanes 5 and 6). These results clearly show the accumulation of deamidated HsTPI in tumors and the effect of treatment with rabeprazole on this protein. The graph (right) shows the significantly increased production of AGEs in tumors treated with rabeprazole compared to untreated tumors. Differences among groups were assessed with one-way ANOVA and Tukey’s test with *p-value* = 0.001 ***.

### Caspase-1 inhibition is related to the accumulation of cellular deamidated HsTPI

To gain more insight in the reason for higher susceptibility of cancer cells for drugs targeting the highly expressed deamidated HsTPI we studied the involvement of caspase-1. HsTPI and some other glycolytic enzymes are substrates of the protease caspase-1[30]. According to relative caspase-1 cleavage specificity, the deamidated sequence of HsTPI (X-Asp-Gly-X) is 152 times more prone to be cleaved by Caspase-1 than its nondeamidated counterpart (X-Asn-Gly-X)[31, 32]. In addition, caspase-1 mRNA expression was found to be significantly decreased in the breast cancer tissues of patients, and treatment with a caspase-1 inhibitor markedly increased the proliferative and invasive abilities of MDA-MB-231 cells[33]. The keystone of our hypothesis implicates the accumulation of deamidated HsTPI in cancer cells. Therefore, we wondered whether caspase-1 plays a role on the absence of deamidated HsTPI in normal cells in contrast with its accumulation in cancer cells. Remarkably, normal cells treated with a caspase-1 inhibitor showed the *de novo* accumulation of acidic isoforms similar to those previously found in cancer cells (Fig. 6a, lanes 4 and 5). Additionally, caspase-1 activity was inhibited by 77%, whereas HsTPI activity was increased by 45% in the noncancer cells (Suppl. Table 3). Moreover, the acidic isoforms of HsTPI from these cells not only adopted the behavior of their counterparts in cancer cells but also had become sensitized to rabeprazole and auranofin (Fig. 6a, upper and bottom, lanes 6 and 7). These normal cells with inhibited caspase-1 activity underwent the aforementioned changes and significantly increased their capacity to produce advanced glycation end products (AGEs) when they were treated with rabeprazole or auranofin (Fig. 6b). Based on these variables, these results show how normal cells can be guided toward “cancer-like” behavior regarding glycolysis, showing characteristic HsTPI electrophoretic profiles, and leading to selective cell apoptosis (Fig. 6 c). Altogether, the results clarify the mechanisms involved in natural accumulation of deamidated HsTPI in cells, leading to the generation of a selective target.

**Figure. 6.**
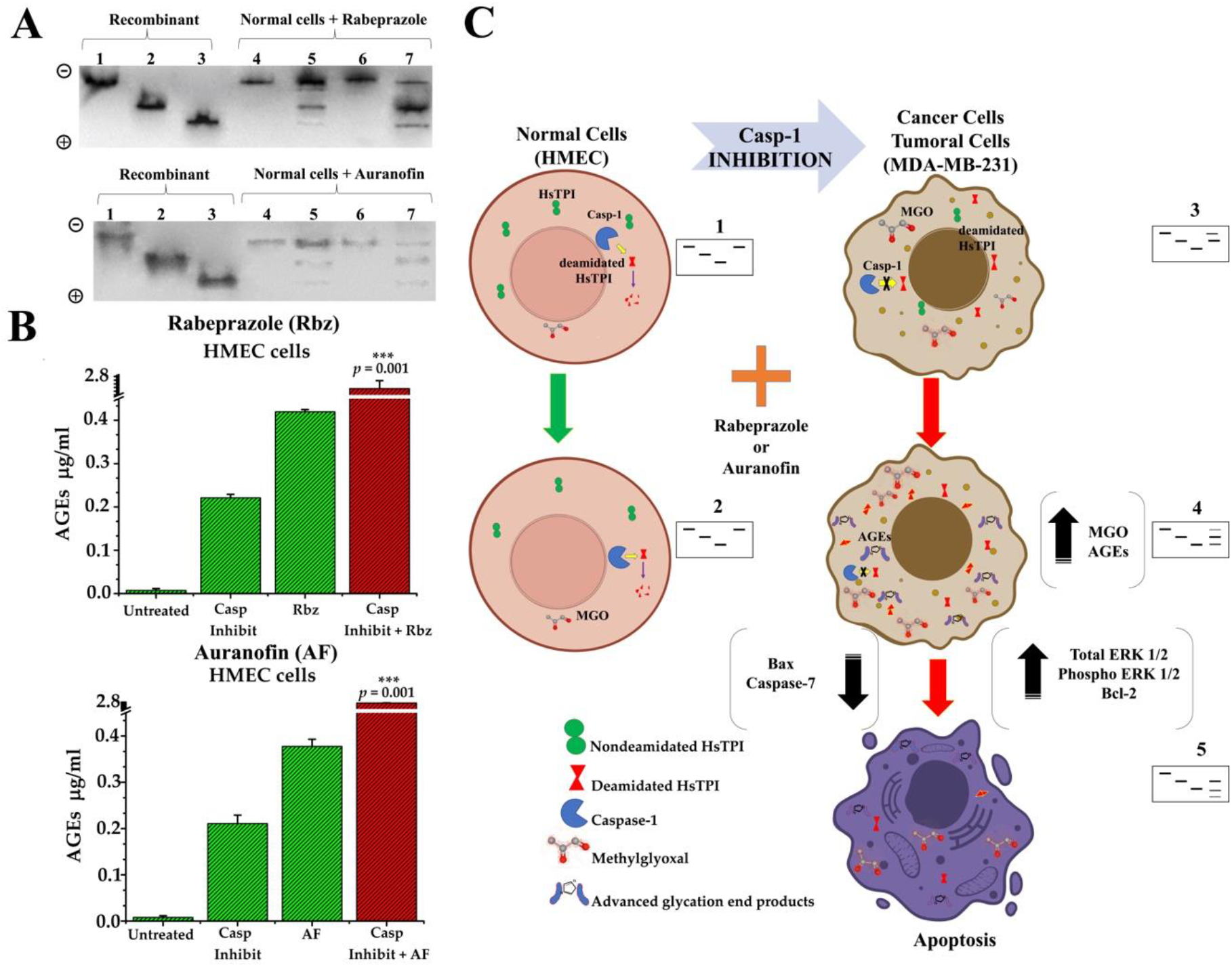
Caspase-1 inhibition in normal cells mimics cancer cells phenotype. (**A**) nPAGE and western blot analysis of HMECs with or without treatment with a caspase-1 inhibitor. Lanes 1 to 3 (10 µg/lane): recombinant nondeamidated, once deamidated, and double deamidated HsTPI, respectively. Lanes 4 to 7 (100 µg protein/lane): proteins from the cellular extract of normal cells (HMECs). Lane 4: cells that were untreated with the caspase-1 inhibitor (top, rabeprazole and bottom, auranofin); lane 5 (top and bottom): cells treated with the caspase-1 inhibitor; lane 6: cells that were treated with rabeprazole (top) and auranofin (bottom), lane 7: cells that were pretreated with the caspase-1 inhibitor and treated with rabeprazole (top) and auranofin (bottom). (**B**) AGEs production in normal cells (HMEC) without the caspase-1 inhibitor, cells that were treated with the caspase-1 inhibitor (Casp Inhibit), and cells that were pretreated with the caspase-1 inhibitor and treated with rabeprazole (upper) and auranofin (bottom). Significant differences were obtained for cells pretreated with the caspase-1 inhibitor and treated with either rabeprazole or auranofin, *p* = 0.001***. (**C**) Schematic illustration of the role of deamidated HsTPI and caspase-1 in normal and cancerous phenotypes (*i*.*e*., MDA-MB-231 cells and tumors). 1, 2, 3, 4, and 5 represent the corresponding nPAGE profiles of HsTPI acidic isoforms from each cell phenotype.

## DISCUSSION

Here we report the selective effect of rabeprazole and auranofin on HsTPI *in vitro, in situ*, and *in vivo*. The derivatization of Cys residues guides the effects exerted by these drugs. Cancer targets are usually selected based on their enhanced expression or even their overactivity [34, 35]. On this basis, human TIM fulfills one requirement for use as a drug target as it is upregulated in cancer cells [4, 36]. Nevertheless, this protein is present in all cells regardless of whether they are cancerous, which significantly diminishes the chosen drug’s cellular selectivity. To address this issue, we propose a new approach to enhancing drug selectivity based on the distinctive characteristics of deamidated HsTPI. First, the presence of deamidated HsTPI in cancer cells but not their normal counterparts. Second, deamidated HsTPI accumulates in cancer cells. Third, deamidated HsTPI is involved in cell-specific cell death induced by rabeprazole and auranofin.

HsTPI is a potential target in cancer mainly by using inhibitors directed to its active site or factors that diminish its expression [37]. Nonetheless, these proposals’ main problem is that the molecular elements involved in the likely therapeutic strategies are present at both pathologic and normal stages. Consequently, an essential disadvantage of the latter approach is that it vastly reduces the safety of such methods. Herein, we have demonstrated that deamidated HsTPI is a distinctive and conspicuous target in triple-negative breast cancer cells but not in their noncancerous counterparts. Hence, our proposal would help the drug design pipeline in the pharmaceutical industry and increase clinical trials’ success rate. MDA-MB-231 cells are triple-hormone receptor-negative breast cancer cells characterized by the minimal expression of estrogen and progesterone receptors and absence of human epidermal growth factor receptor-2 (HER2). Because these cancer cells lack HER2, hormone therapy and drugs targeting HER2 are not helpful, leaving chemotherapy as the main systemic treatment option for triple-negative breast cancer. The primary therapies used for this cancer combine the drugs adriamycin and cyclophosphamide; adriamycin, cyclophosphamide, and paclitaxel; and docetaxel and cyclophosphamide [38]. Since these drugs are directed to targets (mainly nucleic acids and the replication machinery) present in both cancer and normal cells, the side effects of such treatments are considerable and usually severe.

Additionally, their economic burden has a significant impact on patients suffering from this disease [39]. In contrast, our strategy would represent a safer and less expensive therapy for this kind of cancer.

It is important to note that the two different drugs used yielded the same output, especially that caused by the effects on deamidated HsTPI. Accordingly, both treatments exert selective cellular death, mainly attributable to the presence of deamidated HsTPI and its impact on these cancer cells. Consequently, glycolytic flux diminishes, and the concentrations of MGO and AGEs concomitantly increase. Although energy metabolism was affected, the leading cause of selective cell death was the toxic effect of MGO overproduction, as shown in other diseases [40, 41]. Emerging evidence indicates that MGO and AGEs can induce the apoptosis process [42, 43].

Since the response of cancer cells in tumors is affected by factors such as their heterogeneity, components of the microenvironment, and anatomical structures for proper growth [44], the xenograft model was of utmost importance. In this preclinical scenario, the protein concerned was corroborated by the accumulation of acidic isoforms equivalent to deamidated HsTPI in all tumors. Collectively, these results in tumors faithfully mimicked the behavior of HsTPI demonstrated in cell lines. Other proton pump inhibitors (PPIs) have been reported to act as tumor suppressors [6, 45], with proteins upregulated in cancer stages acting as possible targets; however, their most significant weakness is that the proposed targets are found in cancer and normal cells.

The role of deamidated HsTPI as a triple-negative breast cancer-specific target was supported by the change in phenotype of normal cells (HMECs) to a cancer phenotype, based on the described HsTPI behavior. This was achieved by inhibiting cellular caspase-1 activity in HMECs. Most importantly, these cells were sensitized to the drug treatments. Indeed, we previously demonstrated in a cell model that the presence of deamidated HsTPI alone caused sensitization to another PPI, impairing cell growth[10].

The strategy stated here highlights the posttranslational modifications favored in cancer cells, which naturally generate cancer-specific targets. Based on the knowledge acquired herein, pathologic cells should contain a collection of proteins with special skills similar to those shown by HsTPI. That is, structural signatures that usually promote changes in proteins that drive their dead-end elimination (i.e., protein turnover) in normal cells but translate into new functions that lead to adaptive advantages in pathological cells. MGO is a hormetic metabolite related to tumor growth and metastasis [46, 47]. In fact, in this effect, we found a hidden opportunity to push the triple-negative cancer cells to overproduce MGO and commit them to suicide (apoptosis), achieving a novelty and safer strategy on the therapeutics for such highly aggressive cancer. In contrast, cell death would not be as specific for malignant cells if toxic metabolites were administered exogenously. Other types of cancer or even to other pathologies could also adapt our approach for different proteins that meet the characteristics described herein.

According to our findings, three main hallmarks might guide the search for other types of cancers susceptible to being treated using the strategy studied herein—the increase in glycolytic flux, the upregulated HsTPI, and downregulated Casp-1. The presence of all these factors, the coexistence of two of them, or the presence of one of them could indicate the natural accumulation of deamidated HsTPI. In this regard, many cancers might be candidates since they show upregulation of HsTPI (for a detailed list, see [48]). Indeed, the HsTPI upregulation can provide non-deamidated and deamidated HsTPI to the cells as we observed in MDA-MB-231, which is likely reported by others as two types of HsTPIs when they analyzed the proteome of lung squamous carcinoma [49].

Interestingly, monocytic lineage of acute leukemia was previously reported with highly expressed HsTPI [50], and our preliminary findings working with acute leukemia cells show similar results as those in triple-negative breast cancer. We are still working on this to better understand and translate our strategy to a clinical trial shortly.

## Methods

### Reagent and general materials

Luria-Bertani (LB) medium and isopropyl-β-D-thiogalactopyranoside (IPTG) were purchased from VWR Life Science Products (Radnor, Pennsylvania, USA). Glycerol-3-phosphate dehydrogenase (α-GDH), L-lactate dehydrogenase from rabbit muscle, and reduced nicotinamide adenine dinucleotide (NADH) were obtained from Roche (Penzberg, Upper Bavaria, Germany). Immobilized metal affinity chromatography (IMAC) resin was obtained from Bio-Rad (Hercules, California, USA). Sephadex G-25 Fine Resin was obtained from Amersham Biosciences (Amersham, UK). Amicon Ultra 30 kDa filters were purchased from Merck-Millipore Corporation (Billerica, Massachusetts, USA). Fetal bovine serum (FBS), penicillin, streptomycin and trypsin EDTA solutions were purchased from Invitrogen (Carlsbad, USA). The other reagents mentioned were acquired from Sigma-Aldrich (St. Louis, MO, USA).

### Molecular docking studies of deamidated and nondeamidated HsTPI

To carry out the docking studies, the crystallographic structures of HsTPI WT (nondeamidated) and HsTPI N16D (deamidated) that had been deposited in the Protein Data Bank (PDB) were downloaded. Atomic coordinates of the nondeamidated and deamidated proteins (PDB IDs: 2JK2 and 4UNK, respectively) were prepared by removing all water molecules and heteroatoms with PyMOL version 2.0.7 (Schrödinger Inc, NY, USA). The structures were energy minimized with Chimera software [51], and the new coordinates were used for docking calculations. Structures of the drugs rabeprazole and auranofin were obtained from the PubChem Compound Database (https://pubchem.ncbi.nlm.nih.gov) and energy minimized with Avogadro version 1.2. Structures of the nondeamidated and deamidated HsTPIs were prepared by adding hydrogen atoms and Kollman charges (6.00 and 3.999, respectively) with AutoDock Tools (ADT) version 1.5.6 [52]. The molecular docking program AutoDock Vina version 1.1.2 [53] was used with the default settings, and the output files were saved in pdbqt format. Protein receptors and ligands were converted into pdbqt format. The ligand-binding site was defined as the interface of the dimer. After docking, close interactions for binding of the target with the ligands were analyzed and visualized using ADT and PyMOL version 2.0.7.

### Expression and purification of recombinant enzymes

The genes encoding wild-type and mutant (WT, N16D and N16D/N72D) HsTPI were cloned into the vector pET3a-HisTEV as previously reported [12]. The plasmid provides six histidine residues (His6) at the N-terminus of the protein and a tobacco etch virus protease (TEVp) recognition sequence that facilitates protein purification. The plasmids containing inserts (pET3a-HisTEV-wt-HsTPI, pET3a-HisTEV-N16D-HsTPI and pET3a-HisTEV-N16D/N72D-HsTPI) were used to transform the *Escherichia coli* BL21-CodonPlus-RIL strain. The overexpression and purification of recombinants WT (nondeamidated HsTPI), N16D HsTPI (deamidated HsTPI) and N16D/N72D HsTPI (double deamidated HsTPI) were carried out as previously described [12]. Purified proteins were ultrafiltered with Centricon filters (cutoff of 30 kDa for WT and 10 kDa for N16D and N16D/N72D) until the volume reached 0.5 mL, which was repeated 3 times after the addition of 5 mL of buffer containing 100 mM triethanolamine and 10 mM EDTA (pH 7.4) (TE buffer). Finally, the concentrated proteins were precipitated with ammonium sulfate at 75% saturation and maintained at 4 °C. To remove the His_6_-TEV tag, the protein suspension was centrifuged at 12,000 rpm for 20 min at 4 °C, and the pellet was resuspended in 50 mM Tris (pH 8.0), 0.5 mM EDTA and incubated at room temperature for 16 h in the presence of the protease TEVp at 1:50 (w/w) (protease/HsTPI) and 1 mM dithiothreitol (DTT). Thereafter, the incubated sample was loaded into a column containing IMAC resin previously equilibrated with 100 mM triethanolamine (pH 7.4). Enzymes without the His_6_-TEV tag were recovered, ultrafiltered, precipitated with ammonium sulfate and stored at 4 °C until use. To remove the precipitating agent, the protein was centrifuged as mentioned and suspended in TE buffer. The protein concentration was calculated spectrophotometrically at 280 nm with an extinction coefficient of ε = 33,460 M^-1^ cm^-1^ [54]. The purity and integrity of the proteins were verified by sodium dodecyl sulfate polyacrylamide gel electrophoresis (16% SDS-PAGE), and proteins were stained with colloidal Coomassie brilliant blue.

Prior to any assays, the recombinant enzymes were equilibrated in TE buffer and incubated in the presence of 5 mM DTT for 30 min at 4 °C. To remove the reducing agent, the protein was spin filtered in a 1-mL column loaded with Sephadex G-25 Fine Resin previously equilibrated with TE buffer, and the protein concentration was estimated by the absorbance at 280 nm.

### Inactivation assays with nondeamidated (WT) and deamidated (N16D) HsTPI treated with rabeprazole and auranofin

Recently, we demonstrated that the PPI omeprazole selectively inactivates deamidated HsTPI [10]. Furthermore, in a previous work, we showed that among commonly used PPIs (benzimidazole derivatives), rabeprazole was the most efficient PPI in inactivating the TIM of *Giardia lamblia* [55]. Therefore, we chose rabeprazole based on the latter finding; additionally, we tested auranofin to compare the effects of rabeprazole with those of a non-PPI drug. A 100 mM stock solution of rabeprazole was prepared; to acid activate it, the solution was solubilized in 20% dimethyl sulfoxide and 5% 0.1 N HCl, incubated for 30 min at room temperature in the dark, and diluted with TE buffer to obtain a 5 mM solution. A 100 mM stock of auranofin was prepared by dissolving auranofin in 100% ethanol, and serial dilutions were made with TE buffer. For the inactivation assays, the recombinant enzymes were incubated at 0.5 mg·mL^-1^ for 2 h at 37 °C in the presence of 0, 10, 25, 50, 100 and 250 µM rabeprazole or auranofin. After incubation, the samples were diluted, and 5 ng·mL^-1^ and 50 ng·mL^-1^ samples of WT (nondeamidated) and N16D (once deamidated), respectively, were taken to measure their enzymatic activity. Enzyme activity was spectrophotometrically measured (Spectrophotometer Cary 50, Agilent Technologies, CA, USA) by following DHAP synthesis with a coupled system that followed the oxidation of NADH at 340 nm [56]. The results represent the mean of four independent experiments and are expressed as the percent activity *versus* drug concentration, with the activity of the enzyme without the drug set at 100%.

### Quantification of free thiols in recombinant enzymes

Because the principal mechanism of action of the drugs employed here (rabeprazole and auranofin) is thought to be the derivatization of Cys residues, the number of derivatized Cys residues in the recombinant enzymes was determined by using Ellman’s reagent (5,5′-dithiobis-(2-nitrobenzoic) acid, DTNB) [57]. To carry out the experiment described above, 0.5 mg·mL^-1^ protein was incubated without or with 250 and 50 µM rabeprazole and auranofin, respectively, for 2 h at 37 °C. After the incubation period, the proteins were extensively washed by ultrafiltration with Centricon filters to eliminate excess drug, and the protein concentration was estimated by determining the absorbance at 280 nm. Next, an aliquot was taken and used to evaluate the residual activity of assayed proteins. The free thiol (Cys) content of the samples was spectrophotometrically quantified as follows: the basal absorbance of 1 mM DTNB and 5% SDS dissolved in TE was measured at 412 nm (ε _412nm_ = 14.1 mM^-1^·cm^-1^), and the increase in absorbance following the addition of 200 µg of protein was monitored. The number of derivatized Cys residues was indirectly calculated by subtracting the number of free Cys residues in the derivatized enzyme (treated with rabeprazole or auranofin) from the number of free Cys residues in the enzyme in the absence of the drugs. The results represent the mean of at least four independent experiments.

### Determination of fluorescence emission spectra of the recombinant enzymes

The intrinsic and extrinsic fluorescence emission spectra of the proteins were measured using an LS55 spectrofluorometer (Perkin Elmer, Waltham MA, USA). WT or N16D HsTPI (0.5 mg·mL^-1^) was incubated for 2 h at 37 °C with or without 250 and 50 µM rabeprazole and auranofin, respectively. Next, the proteins were extensively washed by ultrafiltration with Centricon filters to eliminate excess drug, and their concentration were recalculated by measuring their absorbance at 280 nm. To determine the intrinsic fluorescence, the enzymes (0.5 mg·mL^-1^) were excited at 295 nm, and the fluorescence emission spectra were recorded from 300 to 500 nm. To determine the extrinsic fluorescence, a stock solution of 15 mM ANS dissolved in methanol was prepared. Then, 100 µM ANS was added to the samples (maintained with slow agitation), which were excited at 385 nm, and the fluorescence emission spectra were monitored from 400 to 600 nm. For each sample reading, the background fluorescence was subtracted (buffer with drug, buffer with ANS, or buffer with ANS and rabeprazole or auranofin). Each spectrum is the average of three scans. The results represent the mean of four independent experiments and are expressed as the percent fluorescence intensity *versus* wavelength.

### Cell culture and identification of cellular deamidated HsTPI by the selective cleavage of Asn-Gly

For general cell culture procedures, the MDA-MB-231 (breast cancer cells) cell line was obtained from American Type Culture Collection (ATCC; Rockville, MD, USA) and maintained in DMEM supplemented with 10% FBS and antibiotics (100 U/mL penicillin and 100 mg/mL streptomycin). Acting as noncancerous (control) cells, HMECs were obtained from ATCC and cultured in mammary epithelial cell basal medium (ATCC #PCS-600-030) supplemented with components of the mammary epithelial growth kit (ATCC #PCS-600-040). For all experiments, both cell lines were used from passages 2–5 and cultured at 37 °C in a humidified 5% CO_2_ atmosphere, detached with 0.05% trypsin in 0.53 mM EDTA, incubated for 5 min at 37 °C, centrifuged at 2000 rpm for 5 min and washed three times with phosphate-buffered saline (PBS).

Since the amino acid sequence of HsTPI contains two Asn-Gly pairs, both cell lines were properly maintained and used to determine the deamidation of cellular HsTPI by selective cleavage with hydroxylamine [18]. A validated method for peptide mapping and sequence analysis of Asn-Gly pairs[19], this method is based on placing the proteins under alkaline conditions (*i*.*e*., pH 9.0) to deprotonate the amide group of the Asn residue, which in turn exert a nucleophilic attack on the adjacent Gly residue located in the C-terminus. This results in the formation of succinimide, which is selectively cleaved in the presence of an excess of hydroxylamine. Notably, deamidated proteins contain Asp (or isoAsp) instead of Asn; thus, under the conditions described for this method, succinimide is not formed, preventing cleavage by hydroxylamine [20].

HMECs (1×10^7^) and MDA-MB-231 cells were incubated for 24 h under the conditions mentioned above. After washing, the cells were lysed in ice-cold RIPA lysis buffer containing protease inhibitors (sc-24948, Santa Cruz Biotechnology, Santa Cruz, CA, USA) and centrifuged at 12,000 rpm for 20 min at 4 °C. The supernatant was collected, and the protein content was quantified by Bradford assay. The protein extracts from each culture were immunoprecipitated with anti-human TIM (H-11) (Santa Cruz Biotechnology) for 1 h at 4 °C. After this, protein A/G agarose (Santa Cruz Biotechnology, sc-2003) was added to the mixtures, which were adequately resuspended and incubated overnight at 4 °C. The mixtures were centrifuged at 2000 rpm for 3 min and washed with PBS 7 times. The immunoprecipitation products were washed with 0.2 M glycine-HCl to separate the protein A/G plus protein-antibody-protein complex. To obtain the isolated cellular HsTPI, samples were centrifuged at 2000 rpm for 5 min, immediately after which the supernatant was taken, and the pH was neutralized with Tris buffer (pH 8.0).

Hydroxylamine cleavage of recombinant and cellular HsTPI was performed as follows. WT (nondeamidated), N16D (once deamidated) and N16D/N72D (twice deamidated) recombinant HsTPI enzymes and cellular HsTPI (1 mg/mL) were separately incubated for 8 h at 45 °C with 2 M hydroxylamine-HCl and 1 M sodium carbonate (Na_2_CO_3_), pH 9.0. Immediately after incubation, the samples were loaded on a 16% SDS-PAGE gel. To analyze the patterns of HsTPI hydrolysis, gels were stained with colloidal Coomassie brilliant blue. The assays were carried out in triplicate to guarantee reproducibility.

### Cell line authentication

Cell line authentication was performed by ATCC, sales order SO0359144, FTA Barcode STRA11162. Briefly, seventeen short tandem repeat (STR) loci plus gender determining locus, Amelogenin, were amplified using commercially available PowerPlex^®^ 18D Kit fro Promega. The cell line samples were processed using ABI Prism^®^ 3500xl Genetic Analyzer. Data were analyzed using GeneMapper^®^ ID-X v1.2 software (Applied Biosystems). Appropriate positive and negative controls were run and confirmed for each sample.

### Native gel electrophoresis and western blot analysis of recombinant enzymes, protein cellular extracts, and tumors

The HsTPI isoforms from cellular extracts were identified with nPAGE combined with western blot analysis using the monoclonal antibody anti-TIM (H11). HMECs or MDA-MB-231 cells (1×10^7^) were exposed to 0, 500, 1000 and 1500 µM rabeprazole and 0, 25, 50 and 100 µM auranofin for 24 h under the abovementioned conditions. Cells were lysed in ice-cold RIPA lysis buffer containing protease inhibitors. Protein quantification was performed by the Bradford method. nPAGE gels were loaded with 1 μg/lane recombinant nondeamidated, N16D (once deamidated), and N16D/N72D (twice deamidated) HsTPI proteins and 100 μg/lane cellular protein extracts. nPAGE gels were prepared with 7% polyacrylamide and Tris-glycine buffer at pH 8.5 [58]. Samples were mixed with native buffer and run at 7 mA and 4 °C for 3 h. Proteins were transferred to polyvinylidene difluoride (PVDF) membranes (0.8 mA/cm^2^, 2 h) in 25 mM Tris buffer containing 192 mM glycine and 20% methanol. The membrane was blocked for 1 h with Tris-buffered saline with 0.1% Tween-20 (TBS-T) supplemented with 5% bovine serum albumin (BSA), washed one time with TBS-T and incubated overnight at 4 °C with anti-TIM (H-11) diluted 1:3000 in TBS-T containing 1% BSA and washed three times with the same buffer. An anti-mouse IgG secondary antibody (diluted 1:5000) conjugated with horseradish peroxidase (HRP) was used to reveal the immunoblot bands by chemiluminescence (Clarity Western ECL substrate, Bio-Rad) following the supplier’s instructions. Blot image acquisition was performed using a ChemiDoc XRS+ system (Bio-Rad Laboratories, Inc., USA).

nPAGE and immunoblotting in tumors were performed as follows. Tumors from nude mice were obtained by dissection and immediately frozen with liquid nitrogen. Afterwards, they were macerated with a mortar until pulverized, immediately after which they were resuspended in PBS and treated as described above. All the assays were carried out in triplicate.

### Cell proliferation and enzymatic assays to assess cultured cells under drug treatment

HMECs and MDA-MB-231 cells were exposed to drugs, after which cell proliferation and the inactivation of cellular HsTPI were determined as follows. A total of 1×10^5^ cells/well in a final volume of 250 µL in six-well plates were exposed to 0, 100, 250, 500, 1000 and 1500 µM rabeprazole and 0, 10, 25, 50, 100, 250 and 500 µM auranofin for 24 h under the culture conditions mentioned above. The cells were detached, washed, and quantified with a hemocytometer. Cell proliferation was measured with 3-(4,5-dimethylthiazol-2-yl)-2,5-diphenyltetrazolium bromide (MTT). HMECs or MDA-MB-231 cells (1×10^3^) were mixed in 100 μL of PBS per well in a 96-well plate, and 10 μL of MTT was added and incubated for 4 h in the dark. The formed formazan crystals were dissolved in DMSO, and the absorbance at 570 nm was measured in an Epoch microplate spectrophotometer (BioTek, Vermont, USA) [59]. The results are shown as the mean of four independent experiments and presented as the percent viability *vs* drug treatment; the viability measured with cellular extracts without treatment was set at 100%. To measure cellular HsTPI enzyme activity, the coupled assay system described above was used. HMECs or MDA-MB-231 cells exposed to drugs were detached, washed and resuspended in TE buffer. Cell suspensions were lysed with 5 freeze/thaw cycles (liquid nitrogen and 40 °C). Then, the protein concentration was determined by Bradford assay. For enzymatic assays, 40-µg protein samples from the cellular extracts were added to 500 µL of enzymatic reaction mixture and spectrophotometrically assessed at 340 nm. The results are presented as the percent activity *vs* drug treatment, and TIM enzyme activity in cellular extracts without treatment was set at 100%.

### L-Lactate measurements

Extracellular L-lactate levels were determined by measuring the reduction of NAD^+^ to NADH by L-lactate dehydrogenase according to Bergmeyer *et al*., 1975 [60]. HMECs and MDA-MB-231 cells (5×10^6^) were treated with 0, 700 and 1000 µM rabeprazole and 0, 70 and 150 µM auranofin for 24 h under the abovementioned culture conditions. Then, the cells were extensively washed with Krebs-Ringer buffer (125 mM NaCl, 5 mM KCl, 25 mM HEPES 1.4 mM CaCl_2_, 1 mM KH_2_PO_4_, 1 mM MgCl_2_, pH 7.4) and resuspended in 250 μL/1×10^6^ cells. The samples were incubated for 10 min at 37 °C under agitation. Then, a 5 mM glucose solution was added and incubated for 0 and 25 min at 37 °C under agitation; at the end of the incubation period, the cells were centrifuged, and the supernatants were carefully taken and stored at -70 °C for the further determination of L-lactate levels. Standard values were determined from a 20 mM L-lactate stock solution in distilled water (standard curve: 0 to 500 μM). To generate the standard curve, aliquots of lactate were added to a cuvette that contained 0.4 M hydrazine, 0.5 M glycine (pH 9.5), 1 mM NAD^+^, and 20 U/mL L-lactate dehydrogenase at 25 °C. The reduction of NAD^+^ was spectrophotometrically (Cary 100 UV/Vis) recorded at 340 nm. To calculate the extracellular lactate level in the samples, 50 μL of supernatant was added to the cuvettes, and NAD^+^ reduction was recorded. Finally, the concentration of lactate was calculated using the extinction coefficient for NADH (ε = 6,200 M^−1^ cm^−1^) and a standard curve. The results are shown as the mean of four independent experiments and expressed as nmol of L-lactate/h/1×10^6^ cells.

### MGO quantification

Intracellular free MGO was spectrophotometrically measured by using 2,4-dinitrophenylhydrazine (DNPH) according to the method of Gilbert and Brandt [61] with modifications [62]. Briefly, 5×10^6^ HMECs and MDA-MB-231 cells were treated with 0, 700, and 1000 μM rabeprazole and 0, 70, and 150 μM auranofin for 24 h under the abovementioned general culture conditions. Cells were resuspended in PBS and lysed with five freeze/thaw cycles, after which 0.45 M perchloric acid was added to each sample, chilled on ice for 10 min, and centrifuged at 12,000 rpm and 4 °C for 10 min. The supernatant was collected and stored at -70 °C for further measurement. Before determining the MGO concentration in the samples, standard MGO values were calculated. Stock solutions of 20 mM DNPH in HCl-ethanol (12:88) and 0.1 mM MGO in distilled water were prepared. Increasing concentrations of MGO (0 to 10 μM) were incubated with 0.2 mM DNPH at 42 °C for 45 min; then, the samples were cooled for 5 min at room temperature, and the absorbance of MGO-bis-2,4-dinitrophenylhydrazone was recorded at 432 nm on a microplate spectrophotometer (Epoch, BioTek, Winooski, VT, USA). Finally, the cell supernatants were taken and used to quantify MGO levels with DNPH. The MGO concentrations from the cells and standard curve were estimated using the extinction coefficient ε = 33,600 M^−1^ cm^−1^ for MGO-bis-2,4-dinitrophenyl-hydrazone and the standard curve. The assays were carried out in triplicate to guarantee reproducibility. The results are expressed as [μM] MGO/1×10^6^ cells.

### AGE quantification

AGEs were determined by using an AGE ELISA kit with the manufacturer’s instructions (MyBioSource, San Diego, CA, USA). HMECs and MDA-MB-231 cells (5×10^6^) were treated with 0, 700, and 1000 μM rabeprazole and 0, 70, and 150 μM auranofin for 24 h under the abovementioned culture conditions. After washing, cells were lysed with RIPA buffer containing protease inhibitors, and the protein extract was separated. The protein concentration was determined with the Bradford method.

The protein concentration in the samples was adjusted to 1 mg/mL, after which the samples were diluted 1:100 and loaded on ELISA plates to determine the AGE concentration. Then, avidin-peroxidase conjugates were added to ELISA wells, and 3,3’,5,5’-tetramethylbenzidine (TMB) was used as the substrate for coloring after the reactant was thoroughly washed out with PBS. Additionally, a standard curve was made with the AGE standard included in the kit. Standard concentrations were 0, 3.12, 6.25, 12.5, 25, 50, 100 and 200 ng/mL. The absorbance at 450 nm was measured within the first 10 min in an Epoch microplate spectrophotometer (BioTek, Vermont, USA). The results are the mean of four independent experiments and are expressed as μg of AGEs/mL.

### Determining the factors related to induced cell death

To assess whether the induced cell death in breast cancer cells was due to apoptosis, assays were conducted as follows. MDA-MB-231 cells (1×10^7^) were treated with 0, 500, 1000 and 1500 μM rabeprazole and 0, 10, 25 and 50 μM auranofin for 24 h. Then, the cells were resuspended in PBS and lysed in ice-cold RIPA buffer with protease inhibitors, and the lysates were stored at -70 °C until use. Protein samples were loaded into 16% SDS-PAGE gels and transferred to PVDF membranes (0.8 mA/cm^2^, 2 h) in 25 mM Tris buffer, 192 mM glycine and 20% methanol. After this, nonspecific binding sites were blocked with TBS-T and 5% BSA, and the membranes were washed one time with TBS-T and incubated overnight at 4 °C with primary antibodies directed against ERK 1/2 (C-9), p-ERK1/2 (12D4), Caspase-7 (10-1-62), Bcl-2 (C-2), Bax (B-9) and β-Actin (C-2) (Santa Cruz Biotechnology, Santa Cruz, CA, USA) at a dilution of 1:1000 in 0.1% TBS-Tween-20 and 1% BSA and washed three times with the same buffer. Proteins were immunoprobed using an HRP-conjugated secondary antibody (dilution 1:3000) and chemiluminescent substrate (Clarity Western ECL substrate, Bio-Rad, Hercules, CA, USA). Blot image acquisition was performed using a Molecular Imager® Gel Doc™ XR+ system (Bio-Rad, CA, USA). The optical density of the protein bands was calculated after background subtraction and normalization to β-Actin using Image Studio 4.0 software (LI-COR Biotechnology).

### TUNEL assays

To further confirm the induced apoptosis, the TUNEL method was employed with an *In situ* Cell Death Detection Kit, Fluorescein 11684795910 (Roche, USA). Briefly, 5×10^4^ MDA-MB-231 cells were grown over glass coverslips into six-well cell culture clusters (Costar, USA) and incubated in the absence or presence of 700 μM rabeprazole or 70 μM auranofin for 24 h at 37 °C. Then, the cells were washed three times with PBS and incubated with Hanks balanced salt solution for 10 min at room temperature. Cells were fixed with 100% methanol at -20 °C for 15 min; finally, the methanol was removed, and the cells were rehydrated by washing with PBS 2 times for 5 min each. Samples were stained with 1 mg/mL 4′,6-diamidino-2-phenylindole (DAPI), washed with PBS and incubated with proteinase K (20 mg/mL) for 30 min at room temperature. Then, the cells were washed with PBS and incubated for 30 min at room temperature with 0.3% H_2_O_2_ in methanol. Immediately after incubation with permeabilization solution (0.1% Triton X-100 in PBS) for 1 h at 4 °C, the cells were washed three times with PBS and incubated in the dark for 1 h at 37 °C with solutions A and B. Analysis was performed by fluorescence microscopy (Axiovert, Carl Zeiss. Germany).

### Caspase-1 assays

Because HsTPI accumulated in breast cancer cells but not in HMECs, we hypothesized that this condition is important to generate a favorable response to drug treatments. Thus, we performed assays to mimic HsTPI accumulation in normal cells by inhibiting caspase-1 as follows.

A stock solution (50 mM) of the selective caspase-1 inhibitor VX-765 [63], from BioVision (Milpitas, CA) was prepared in DMSO and diluted in HMEC medium to the appropriate concentrations. HMECs (5×10^6^) were incubated for 24 h as previously mentioned with or without 500 µM caspase-1 inhibitor. Then, 0.5 and 1.5 mM auranofin and rabeprazole, respectively, were added to the samples and incubated for another 24 h under the same conditions. At the end of incubation time, the cells were detached, washed and prepared for the following assays as previously described: nPAGE, cell viability assay, HsTPI activity assay, AGE quantification, assays to assess factors related to the induced cell death, and TUNEL assays. To corroborate the inhibition of caspase-1, caspase-1 activity was measured. In a buffer containing 100 mM HEPES (pH 7.2), 100 mM NaCl, 0.2% CHAPS, 20 mM EDTA, 10% glycerol and 10 mM DTT, 500 μg of total extract of cellular proteins, the reaction was initiated in a 96-well plate (total reaction volume was 100 µL/well) by the addition of 200 µM Ac-YVAD-pNA substrate.

The plate was then incubated for 4 h at 37 °C, and at the end of the incubation period, the absorbance at 405 nm was read using a microplate ELISA reader.

This assay is based on the ability of the active enzyme to cleave the chromophore pNA, as measured through determining the absorbance at 405 nm.

### *In vivo* experiments

Five-to seven-week-old female athymic Balb/c nude mice (Nu/Nu) were obtained from the Círculo ADN S.A. de C.V. and housed in the Unidad de Investigación Preclínica of the National Autonomous University of Mexico under specific pathogen-free (SPF) conditions with free access to autoclaved food and water; the mice were cared for following NIH guidelines for laboratory animals. For studies of the presence and effect of deamidated HsTPI in tumors, the group sizes were chosen based on previous experience, and mice were randomly assigned to one of the following experimental groups: nontreatment (*n*=4), subcutaneously injected once with 20×10^6^ MDA-MB-231 cells and intraperitoneally administered sterile PBS three times a week; treated (*n*=6), subcutaneously injected once with 20×10^6^ MDA-MB-231 cells and intraperitoneally administered 50 mg/kg rabeprazole three times a week; pretreated (*n*=2), subcutaneously injected once with 20×10^6^ MDA-MB-231 cells previously incubated with 1 mM rabeprazole for 24 h. The viability of the cells was verified before injection, and the injection volume was adjusted to guarantee that at least 20×10^6^ of the injected cells were alive. Human cancer cells were implanted in mice by subcutaneous injection in the dorsal flank. The number of animals and all procedures followed protocols approved by the Institutional Animal Care and Use Committee for Care and Use (IACCUAC protocol UNIPREC-19-020).

Rabeprazole was intraperitoneal applied 7 days after the implantation of MDA-MB-231 cells. The animals were weighed, and the tumor length and width were measured using a digital caliper three times a week for 34 days. Using an established formula (0.52 × (length of the ‘long axis’ of the tumor) × (length of the ‘short axis’ of the tumor) ^2^, tumor sizes were converted into tumor volumes [64].

The experiment finished 24 h after the last drug application in each group, after which the animals were killed in a CO_2_ chamber. Tumors (when they were present) were dissected and prepared to analyze HsTPI as mentioned above.

## Acknowledgments

We thank to Bruno López-Fuentes for critical revision of the language. Daniela Kayani Rangel-López designed Figure 6C. This research was funded by the Recursos Fiscales para Investigación Program from the Instituto Nacional de Pediatría, S.S. Grant Number 2019/072.

